# Replisome preservation by a single-stranded DNA gate in the CMG helicase

**DOI:** 10.1101/368472

**Authors:** Michael R. Wasserman, Grant D. Schauer, Michael E. O’Donnell, Shixin Liu

## Abstract

The eukaryotic replicative helicase CMG is assembled at replication origins and is thought to remain topologically closed until termination. Upon encountering a lesion, CMG must vacate a stalled fork to allow DNA repair. However, the fate of CMG under these stress conditions remains unclear. Here, using correlative single-molecule fluorescence and force microscopy, we show that when uncoupled from a DNA polymerase, CMG opens a single-stranded (ss) DNA gate to traverse a forked junction and reside on double-stranded (ds) DNA. Surprisingly, CMG undergoes rapid diffusion on dsDNA and can transition back onto ssDNA for continued fork progression. The accessory protein Mcm10 is required for robust ssDNA gating. These results reveal an Mcm10-induced pathway that preserves CMG on DNA and allows it to access a repaired fork for swift replication recovery.

---

The eukaryotic replicative helicase CMG (Cdc45-Mcm-GINS) is assembled and activated at origins of replication by numerous factors (Bell and Labib, 2016; Deegan and Diffley, 2016). A mature CMG consists of the Mcm2-7 hexameric ATPase forming a two-tiered ring and the accessory subunits Cdc45 and GINS bracing the N-terminal tier of the ring (Costa et al., 2011; Yuan et al., 2016). After origin firing, CMG unwinds dsDNA by tracking along one strand in the 3’-5’ direction while excluding the other strand from its central channel (Fu et al., 2011; O’Donnell and Li, 2018). During normal synthesis, CMG forms a complex with the leading-strand DNA polymerase (Pol ε) (Langston et al., 2014). Upon encountering a DNA lesion, CMG uncouples from the stalled Pol ε and needs to vacate the replication fork so repair factors can gain access (Berti and Vindigni, 2016). However, it remains unclear whether a repaired fork can be restarted by the same CMG that is retained on DNA, or if replication recovery relies on a distant replisome traveling from another origin.

To gain a better understanding of CMG dynamics, we prepared *Saccharomyces cerevisiae* CMG labeled with a Cy3 fluorophore on Cdc45 (Figure S1) and combined single-molecule fluorescence microscopy and optical tweezers (Hashemi Shabestari et al., 2017) to directly observe the behavior of CMG on DNA in real time (Figure 1A). We first examined CMG on a ssDNA substrate prepared from biotinylated phage λ genomic dsDNA [48.5 kilobase pairs (kbp) in length] and tethered between two optically trapped beads (Figure S2). We found that, when supplemented with Mcm10—an essential eukaryotic replisome protein (Baxley and Bielinsky, 2017; Bell and Labib, 2016)—CMG can readily bind to ssDNA and exhibit unidirectional translocation (Figure 1B). The rate of translocation was measured to be 10.4 ± 0.9 nucleotides per second (nt/s) (mean ± SEM; *N* = 62) with 1 mM ATP at room temperature (Figure 1C). CMG was observed to be immobile in the absence of ATP (Figure S3). Such ATP- dependent directed movement most likely requires strand encirclement via interactions between DNA and the central pore of the Mcm2-7 ring, as supported by structural studies of ring helicases/translocases from the same ATPase family (O’Shea and Berger, 2014). Moreover, loaded CMG can withstand high salt (0.5 M NaCl) and high tether tension [>80 picoNewton (pN)] (Figure S4), further suggesting that it is encircling ssDNA. These results provide direct evidence that CMG can load onto ssDNA without a free end and translocate in a directional and processive manner.

**Figure 1.**
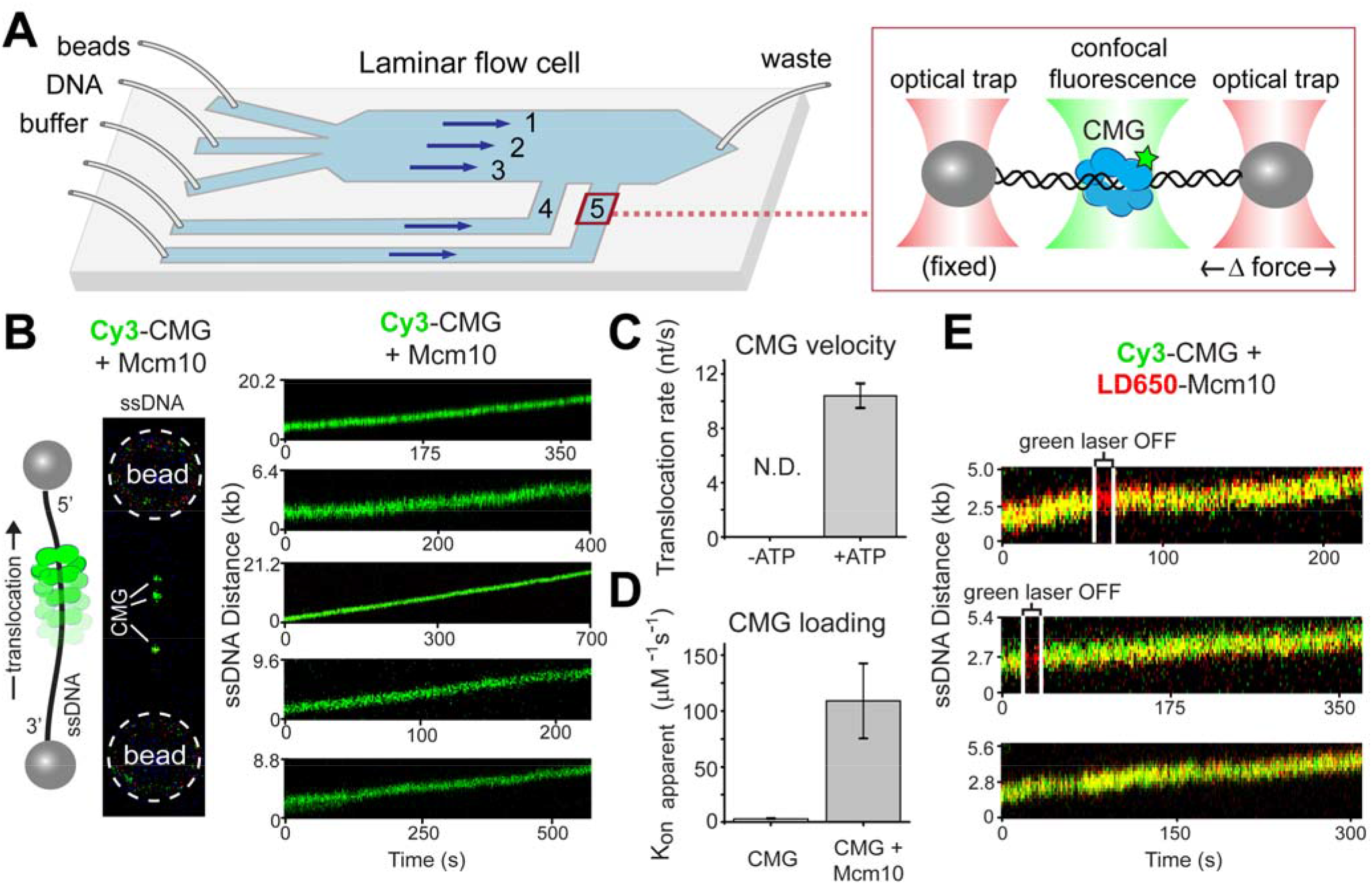
CMG undergoes directional translocation on ssDNA. (**A**) Schematic of the experimental setup. The illustration in the zoom-in box was not drawn to scale. (**B**) (Left) Cartoon and 2D scan of a tethered ssDNA loaded with multiple Cy3-CMGs. (Right) Representative kymographs of CMG movement in the presence of 1 mM ATP and 10 nM Mcm10. (**C**) Comparison of CMG translocation rate on ssDNA in the absence and presence of ATP (*N* = 43 and 62 CMGs, respectively). (**D**) Comparison of CMG loading efficiency in the absence and presence of Mcm10 (*N* = 6 and 9 tethers, respectively; *P* < 0.03). (**E**) Representative kymographs demonstrating co-migration of CMG (green) and Mcm10 (red) on ssDNA. The green laser was occasionally turned off to confirm the fluorescence signal from Mcm10. Error bars in panels C and D represent standard error of the mean (SEM) generated via bootstrapping.

Next we assessed the role of Mcm10 in CMG interaction with ssDNA. We found that omission of Mcm10 decreased the loading efficiency of CMG on ssDNA by ∼65-fold (Figure 1D), suggesting that Mcm10 greatly enhances CMG’s ability to open its ring and encircle ssDNA. Consistent with this picture, using Mcm10 labeled with an LD650 fluorophore and two excitation lasers (532 nm and 638 nm) to detect CMG and Mcm10 fluorescence signals respectively, we showed that CMG and Mcm10 co-localize and co-migrate on ssDNA (Figure 1E). Hence, CMG forms a complex with Mcm10 during ssDNA translocation. We also performed a bulk helicase assay to corroborate the single-molecule data. Without Mcm10, CMG can unwind a ^32^P-oligonucleotide with a 5’ flap annealed to circular M13 ssDNA (Ilves et al., 2010; Langston et al., 2014); however, Mcm10 significantly stimulated unwinding (Figure S5). Due to the 3’-5’ unwinding polarity of CMG (Moyer et al., 2006), the helicase must load and encircle the circular ssDNA in order to unwind this flap. Taken together, these results demonstrate the existence of an Mcm10-stimulated “ssDNA gate” in CMG that enables strand passage. Unless noted otherwise, Mcm10 was thus included throughout this study and we refer to the CMG-Mcm10 complex as CMGM.

We next examined the behavior of CMGM on dsDNA. CMGM displayed minimal affinity to a tethered phage λ dsDNA at low force; by contrast, CMGM binding was readily observed upon application of high tension (>65 pN) to the tether (Figure 2A). We posited that this is due to binding of CMGM to force-induced ssDNA regions. To confirm this interpretation, we used the eukaryotic ssDNA-binding protein RPA labeled with AlexaFluor488 (A488) to mark ssDNA regions of the tether. Areas of RPA fluorescence emerged when high tension was applied to the tether (Figure 2B), indicating that stretches of dsDNA are melted into ssDNA as characterized previously (van Mameren et al., 2009). Interestingly, lowering the tension back to 10 pN led to strand reannealing and ejection of RPA from DNA, which is demonstrated by disappearance of the RPA fluorescence signal (Figure 2B). The observation that RPA rapidly dissociates in favor of DNA rehybridization entails that RPA alone cannot keep two complementary ssDNA strands separated unless the chromosome is under high tension, or at least one strand is occupied with other proteins or nucleic acids—a concept that may have important implications for RPA-ssDNA signaling in the nucleus.

**Figure 2.**
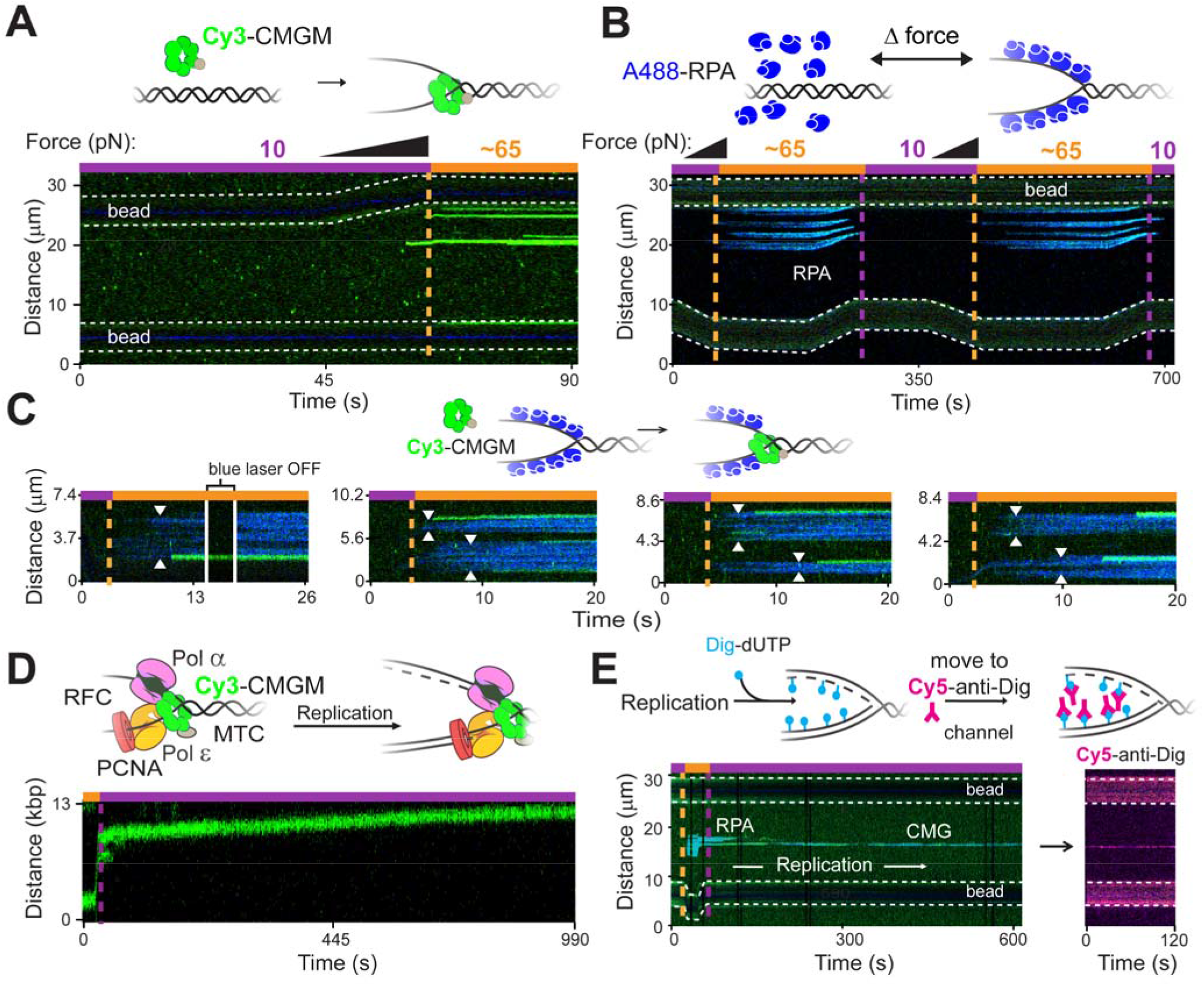
CMGM assembles onto an RPA-coated DNA fork and restarts replication. (**A**) Binding of CMGM (green) to a dsDNA tether is minimal at low force, but greatly enhanced at high force. During the force ramp, one bead held by a movable optical trap is pulled further apart from the other bead held by a fixed trap. (**B**) Formation and collapse of ssDNA regions—marked by fluorescently labeled RPA (blue)— via force manipulation of a dsDNA tether. (**C**) CMGM preferentially loads onto ss-ds fork junctions. Arrows indicate edges of ssDNA regions marked by RPA. (**D**) Representative kymograph showing directed motion of CMGM after binding the fork junction in the presence of unlabeled components required for replication. (**E**) Representative kymograph demonstrating that CMGM loading at the fork leads to active replication. Cy3-CMGM and other unlabeled replisome components were loaded at a DNA fork marked by A488-RPA in the presence of Dig-dUTP. After incubation, the post-replication assembly was moved to a separate channel containing Cy5-anti-Dig for detection of nascent DNA (magenta).

We then used two-color fluorescence detection to directly analyze the co-localization of Cy3-CMGM and A488-RPA. We found that CMGM predominantly (82%, *N* = 71) binds to the edges of RPA-coated ssDNA regions, indicating its preference for ss-ds fork junctions (Figure 2C). If the loaded CMGM encircles ssDNA at the fork junction and functions as a replicative helicase, it is expected to support replisome assembly and fork progression. To test this hypothesis, we complemented Cy3-CMGM with the numerous factors required for replisome-mediated DNA synthesis *in vitro* (Lewis et al., 2017). Force was raised to ∼65 pN to promote CMGM loading at the fork, and then reduced to favor nucleotide addition over exonucleolysis by the polymerase (Wuite et al., 2000). We observed that, upon binding to the fork, CMGM underwent directional movement with a rate of 7.0 ± 1.0 nt/s (mean ± SEM; *N* = 33) at room temperature (Figure 2D), consistent with the replication fork speed measured *in vivo* (Sekedat et al., 2010).

To seek further evidence that CMGM loading at the fork can lead to active replication, we attempted to directly visualize newly synthesized DNA. First we monitored Cy3-CMGM binding to the optically trapped DNA using the same protocol as described above, but with a replication mixture containing digoxigenin-conjugated deoxyuridine triphosphate (Dig-dUTP). After ∼5 minutes of incubation at low force, the tethered complex was moved to a separate channel containing Cy5-labeled anti-digoxigenin antibodies (anti-Dig) (Figure S6A). Nascent DNA stained by Cy5-anti-Dig was indeed observed exclusively at positions where Cy3-CMGM was located (Figures 2E and S6B). We note that Dig-dUTP severely inhibits the rate of synthesis (Figure S6C); thus we did not observe a Cy5-anti-Dig tract length commensurate with the amount of CMGM movement observed without Dig-dUTP. Nonetheless, the observations of CMGM translocation and CMGM-dependent DNA synthesis together strongly suggest that CMGM at the fork is functional for replisome assembly.

Next we sought to follow the fate of CMG once the polymerase becomes uncoupled from the helicase under replication stress. To mimic this situation, we applied the sequential force protocol (high tension followed by low tension) to the dsDNA tether in the presence of only Cy3-CMG and Mcm10 without the other replisome components. Unexpectedly, CMGM was observed to frequently switch to a rapidly diffusive mode upon lowering the force (31%, *N* = 200), traversing up to tens of kbp of dsDNA (Figure 3A). Two-color experiments showed that Mcm10 travels with CMG during diffusion on dsDNA (Figure S7). We interpret this “mode-switch” phenomenon as CMGM transitioning from ssDNA to dsDNA, again necessitating opening of the ssDNA gate in CMG to pass one strand. This interpretation was validated using A488-RPA to distinguish ssDNA from dsDNA. When force was lowered, RPA-bound ssDNA regions reannealed to form duplexes, and CMGM originally residing at the fork junction departed the fork and underwent diffusion (Figure 3B). Conversely, upon re-introduction of ssDNA regions by applying high tension again, the rapidly diffusive CMGM soon located a newly formed ss-ds fork junction, at which diffusion halted (Figure 3C). Furthermore, when moved to another channel containing free replisome factors in solution, a CMGM in the diffusive mode can transition back to the ssDNA-binding mode and recover directed and processive movement with a speed consistent with fork progression (Figure 3D). This result indicates that the same CMGM that has left the fork junction is able to re-enter a fork and nucleate an active replisome. Importantly, in the presence of replisome components (Figures 2D and 3D), CMGM rarely entered the diffusive mode (10%, *N* = 224), indicating that disengagement of CMG from an active DNA polymerase substantially stimulates its transition from ssDNA to dsDNA.

**Figure 3.**
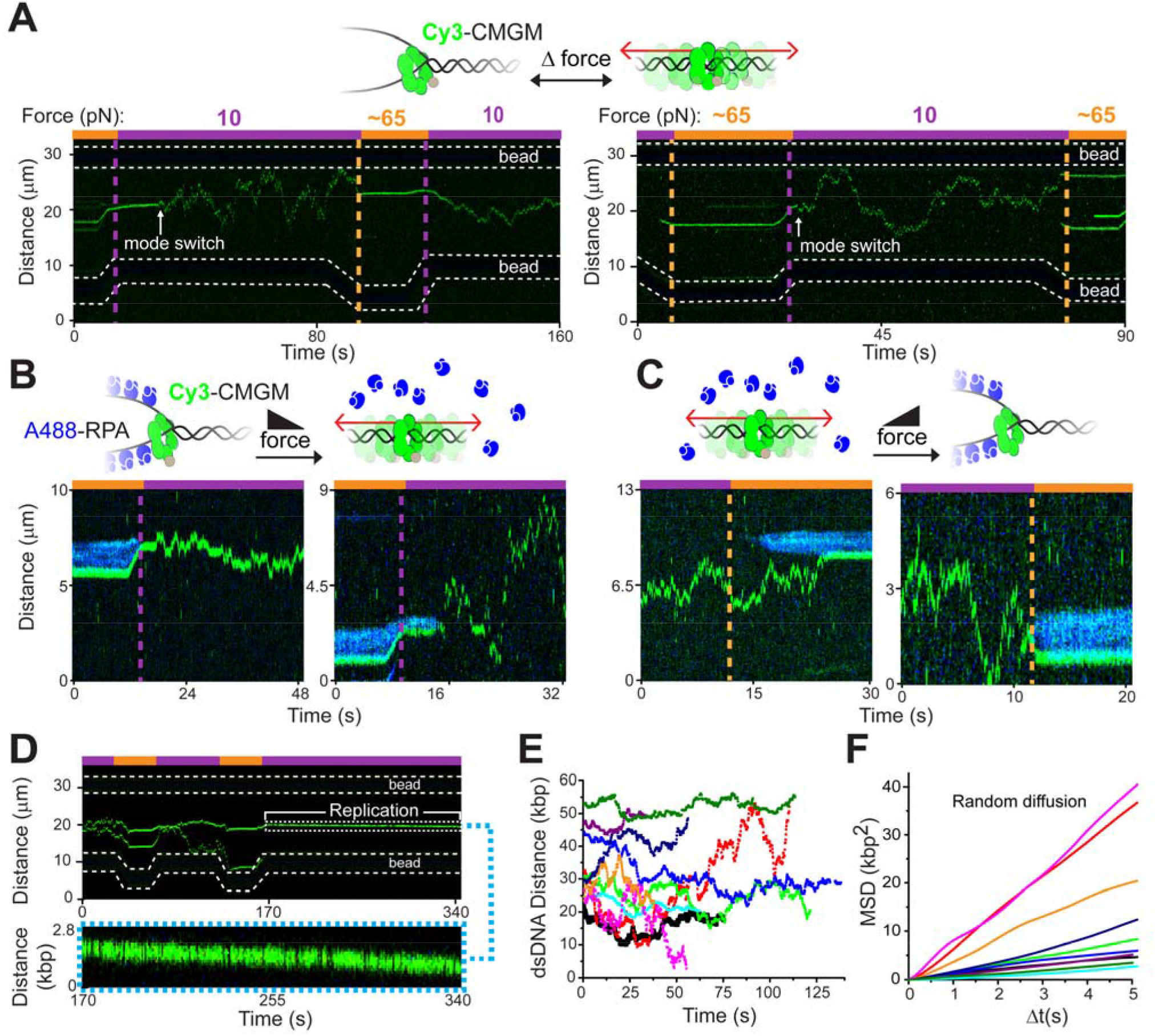
CMGM without a coupled polymerase can jump the fork and diffuse on dsDNA. (**A**) Representative kymographs of CMGM (green) switching between non-diffusive (high force) and diffusive (low force) modes. In the right example, two more CMGMs bound DNA from solution at the end of the kymograph when the tension was raised. (**B**-**C**) Representative kymographs displaying CMGM mode-switching at ss-ds fork junctions. RPA (blue) was used to distinguish ssDNA from dsDNA regions. (**D**) Representative kymograph demonstrating that a diffusive CMGM re-entering a fork in the presence of replisome factors leads to directed translocation. (**E**) Examples of CMGM in the diffusive mode. Trajectories are offset for clarity. (**F**) Mean square displacement (MSD) analysis of the trajectories shown in panel E (color matched). The linear dependence of MSD on Δ*t* suggests random diffusion.

Mean square displacement analysis showed that CMGM motion over long regions of dsDNA is a random walk, with a diffusion coefficient of 1.81 ± 0.48 kbp^2^/s (mean ± SEM; *N* = 35) at room temperature (Figures 3E, 3F, and S8). We presume that the rapidly diffusive form of CMGM is topologically linked to dsDNA, given our observation that diffusion persisted after high-salt wash and application of orthogonal hydrodynamic force (Figure S9), and previous work suggesting that CMG (Langston and O’Donnell, 2017) and Mcm2-7 hexamer (Evrin et al., 2009; Randell et al., 2006; Remus et al., 2009) are able to encircle and slide over dsDNA. Although we cannot rule out the possibility that CMGM can be diffusing on the surface of dsDNA, in either case mode-switching from ssDNA to dsDNA requires the ssDNA gate to open to either accept the complementary strand or to expel the original strand. As with the Mcm10-dependent loading of CMG onto ssDNA (Figure 1D), Mcm10 is also essential for robust transition to the diffusive mode and preservation of CMG on dsDNA: when Mcm10 was omitted from the assay, the vast majority of CMGs (89%, *N* = 140) dissociated from DNA after collapse of the ss-ds fork junction (Figure S10).

To summarize, we show that CMGM displays unexpected structural plasticity that enables it to switch between multiple functional modes. When uncoupled from a polymerase under replication stress, CMGM can transition from ssDNA to dsDNA (Figures 4A and 4B). DNA repair processes can involve fork reversal and recombination intermediates that are incompatible with retaining a replisome at the fork (Berti and Vindigni, 2016; Bhat and Cortez, 2018). Indeed, single-molecule studies in the phage T4 system indicate that the replicative helicase is removed from the fork during reversal (Manosas et al., 2012). Prokaryotic systems have evolved mechanisms of reloading the helicase onto the ssDNA of a collapsed fork for reactivation (Marians, 2018). However, eukaryotes have not been documented to possess soluble CMG helicases for *de novo* reassembly. The ssDNA-to-dsDNA mode-switching we observe for CMGM provides an elegant way of helicase vacating a stalled fork while preserving it on DNA. After the repair is completed, the reverse mode-switch—from dsDNA to ssDNA—allows CMG to re-enter the fork to resume replication without doing so from the solution phase (Figures 4C and 4D). We provide evidence that Mcm10 stimulates these transitions between ss and dsDNA and remains associated with CMG during these ssDNA-gating processes. We refer to this mechanism as “replisome preservation”: while CMGM itself can nucleate a replisome by recruiting soluble components, it is also possible that other CMG-associated factors, or even the entire replisome, stay with CMG during these transitions. When two opposing forks converge upon replication termination or during interstrand crosslink repair, CMG is demonstrated to be targeted by the ubiquitin ligase and segregase for unloading from DNA (Amunugama et al., 2018; Maric et al., 2014). But whether it is converging CMGs, CMG binding to dsDNA, or some other signal that triggers CMG ubiquitylation remains to be established. Moreover, ubiquitylation-dependent CMG disassembly takes over 20 minutes for significant amounts of removal (Dewar et al., 2017), making it unlikely that a rapidly diffusing CMG is disassembled before returning to a fork.

**Figure 4.**
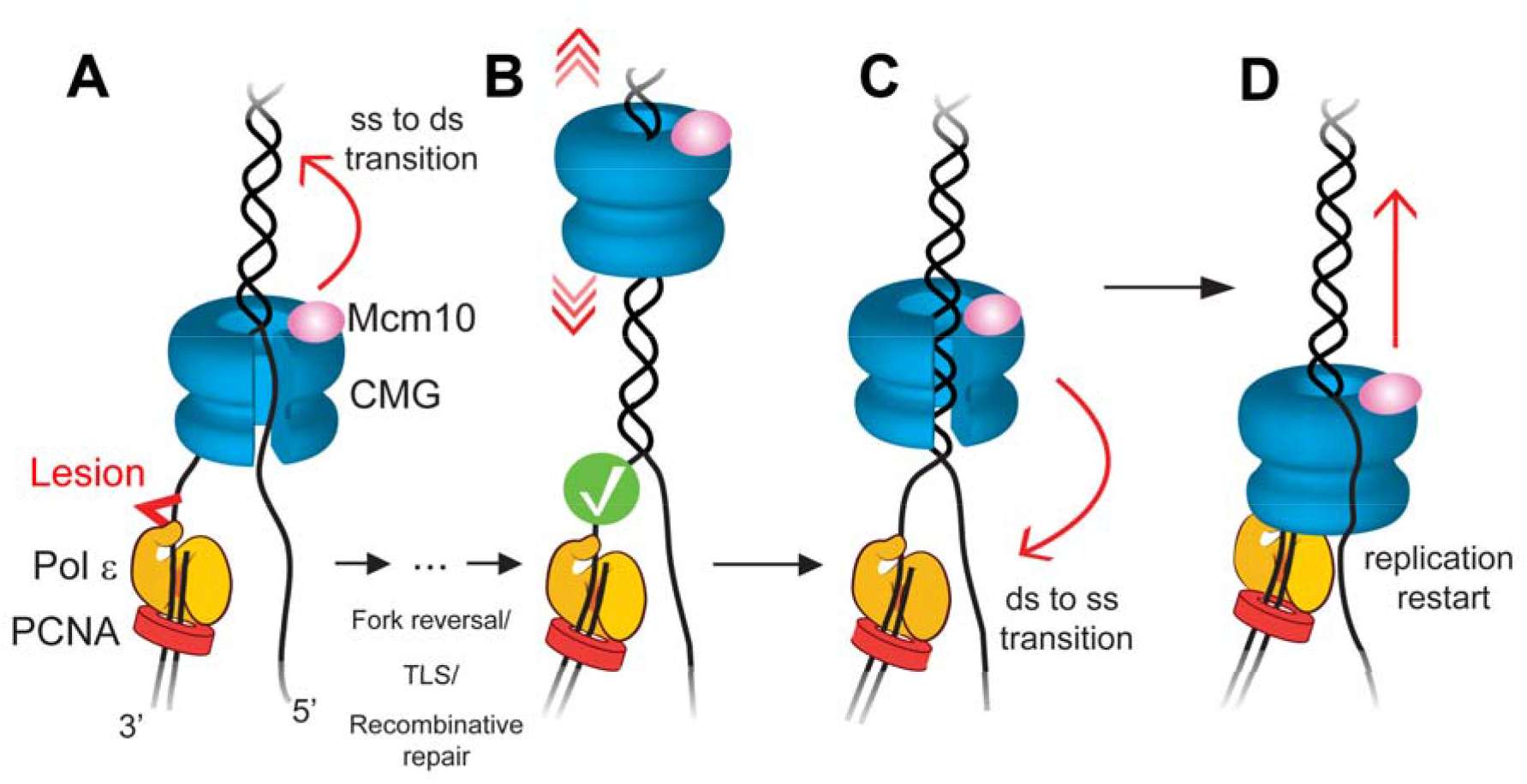
Model for CMG gating and replisome preservation. (**A**) CMG-Mcm10 uncouples from the polymerase and transitions from ssDNA to dsDNA when the replisome stalls during replication stress, for example upon encountering a lesion. (**B**) Upon decoupling, CMGM opens the ssDNA gate and transitions to a fast diffusive mode on dsDNA. Meanwhile, lesions are repaired (check mark) through a multitude of stress response pathways available to the cell such as fork reversal, translesion synthesis (TLS), or recombinative repair. (**C**) When the stress response is complete, CMGM can re-enter the fork by transitioning from dsDNA to ssDNA. (**D**) CMGM nucleates replisome assembly at the restored fork, leading to replication restart.

The transitions between ss and dsDNA binding modes require opening of the Mcm2-7 ring to pass a strand either into the central channel or out of it. This ssDNA gate may also have utility in origin firing. Two CMGs are assembled head-to-head around dsDNA at origins with their N-termini facing each other (Bell and Labib, 2016). Our previous cryo-electron microscopy structure shows that CMG encircles one strand and translocates in an N-first orientation (Georgescu et al., 2017), which is recently supported by work from Diffley and colleagues (Douglas et al., 2018). Thus, at an origin CMG must pass the non-tracking strand to the exterior—possibly through the ssDNA gate documented here—in order for the two head-to-head CMGs to move past one another and establish bidirectional replication. Mcm10, the final origin-firing factor (Bell and Labib, 2016), may directly regulate gate opening and strand passage during this process. The Mcm2/5 interface is used for loading Mcm2-7 around dsDNA during origin licensing (Samel et al., 2014; Ticau et al., 2017); however, this interface is occluded by Cdc45 and GINS in the CMG complex (Costa et al., 2011). Thus, whether another gate in the Mcm ring is employed for ssDNA passage, such as the Mcm4/6 interface (Yuan et al., 2016), requires further study.

## ACKNOWLEDGMENTS

We thank O. Yurieva and D. Zhang for assistance with protein purification, and other members of the O’Donnell and Liu laboratories for discussions. This work was supported by a postdoctoral fellowship from the Anderson Cancer Center at The Rockefeller University (to M.R.W.), the Robertson Foundation, the Quadrivium Foundation, a Monique Weill-Caulier Career Award, a Basil O’Connor Starter Scholar Award from the March of Dimes (#5-FY17–61), and a Kimmel Scholar Award (to S.L.), and National Institutes of Health (T32 CA009673 to G.D.S., R00 GM107365 to S.L., and R01 GM115809 to M.O.D.). M.O.D is a Howard Hughes Medical Institute investigator.

## MATERIALS AND METHODS

### SFP synthase reagents for protein labeling

Site-specific labeling of CMG and Mcm10 used SFP synthase (4’-phosphopantetheinyl transferase), which specifically recognizes a short peptide tag and catalyzes the covalent transfer of CoA-functionalized moieties to a single serine residue within the tag via a phosphopantetheinyl linker. SFP synthase was purified on a nickel-NTA column as previously described (Yin et al., 2006). Cy3 and LD650 [a photostable version of Cy5 (Altman et al., 2011); Lumidyne Technologies] were functionalized by CoA and purified by HPLC as previously described (Yin et al., 2006).

### Preparation of Cy3-CMG

To obtain CMG labeled with a single Cy3, we modified a *S. cerevisiae* strain in which all 11 subunits of CMG have been integrated under inducible control (Georgescu et al., 2014) by inserting the “S6” peptide (GDSLSWLLRLLN) (Zhou et al., 2007) between the C-terminus of Cdc45 and its 3× FLAG tag. To insert the S6 sequence, we transformed and co-expressed Cas9, gRNA targeting a region downstream of the Cdc45 C-terminus, and a ssDNA encoding the S6 peptide into the CMG co-expression strain. Nourseothricin-resistant Cas9-NAT (Addgene #64329) and hygromycin-resistant gRNA-ura-HYB (Addgene #64330) plasmids were gifts from Yong-Su Jin (Zhang et al., 2014). The gRNA sequence of the gRNA-ura-HYB plasmid was mutated with the Q5 mutagenesis kit (New England Biolabs) to ACTAGTTAACAATCCACTCA in order to hybridize with a 20-nt region downstream of Cdc45. The DNA donor (ACGGACACTTACTTGGTTGCTGGGTTAACACCTAGGTATCCTCGCGGACTAGACACGATACACACAAAAAAACCTATTTTGAATAATTTCAGCATGGCGTTTCAACAAATAACTGCAGAAA CGGATGCTAAAGTGAGAATAGATAATTTTGAAAGTTCCATAATTGAAATACGTCGTGAAGA TCTTTCACCATTCCTGGAGAAGCTGACCTTGAGTGGATTGTTAGGATCT<u>GGCGATAGCCT GAGCTGGCTGCTGCGCCTGCTGAAC</u>ACTAGTGACTACAAAGACCATGACGGTGATTATAA AGATCATGACATCGACTACAAGGATGACGATGACAAGTAGGGATCCGGCTGCTAACAAAGCCCGAAAGGAAGCTGAGTTGGCTGCTGCCACCGCTGAGCGGCCGCCAGGTGGATGGGGGTAATATAATTGTATCTATGTATCTGG) was purchased from IDT. The underlined sequence encodes the S6 peptide. Co-transformants of these constructs were grown under auxotrophic (–Ade –His –Leu –Trp –Ura) and antibiotic (hygromycin and nourseothricin) selection in SC glucose, streaked on selective plates, and colonies were sequenced to confirm the existence of the S6 tag. The resultant CMG-S6 construct was overexpressed and purified as previously described (Georgescu et al., 2014). CMG-S6, SFP, and Cy3-CoA were then incubated at a 1:2:5 molar ratio for 1 hour at room temperature in the presence of 10 mM MgCl_2_. Excess dye and SFP were removed by purification on a Sepharose 6 column (GE Healthcare) with a buffer containing 150 mM NaCl, 10% glycerol, 20 mM Tris-HCl pH 7.5, 1 mM MgCl_2_, and 1 mM DTT. The final labeling ratio was estimated to be ∼100% from the extinction coefficients of Cy3 and CMG (Figure S1A). Peak fractions were aliquoted, flash frozen, and stored at –80 °C.

### Preparation of LD650-Mcm10

To obtain Mcm10 labeled with a single LD650, we overexpressed *S. cerevisiae* Mcm10 bearing a hexahistidine tag followed by the S6 peptide at the N-terminus and a 3×FLAG tag at the C-terminus in *Escherichia coli* BL21-DE3 codon plus RIL cells (Stratagene) under ampicillin and chloramphenicol selection. 12 L of cells were grown to an OD_600_ of 0.6 and induced for 4 hours at 20 °C. The cells were lysed by French press and clarified by centrifugation at 50,000× g in the presence of 2 U/mL DNase I (New England Biolabs) and 1 mM PMSF for 30 minutes. Lysate was applied to a column packed with 4 mL of anti-FLAG M2 agarose (Sigma-Aldrich) pre-equilibrated with buffer A (750 mM NaCl, 100 mM potassium glutamate, 30 mM Tris-OAc pH 7.9, 15 mM imidazole, and 10% glycerol). The bound material was washed with a 20× column volume (CV) of buffer A followed by elution with 25 mL of buffer A containing 0.2 mg/mL 3×FLAG peptide (EZ Biolab)—pausing 30 minutes every CV— directly onto a 1-mL Ni-NTA column (GE Healthcare). The Ni-NTA column was subsequently washed with 20× CV buffer A and eluted with a linear 15-700 mM imidazole gradient in buffer A. Mcm10-S6 eluted between 180 and 350 mM imidazole. Peak fractions were pooled and dialyzed against 1 L of 300 mM NaCl, 30 mM Tris-OAc pH 8.0, and 10% glycerol overnight at 4 °C. Mcm10 was subsequently labeled with LD650 at the N-terminus by incubating Mcm10-S6, SFP, and LD650-CoA at a 1:2:3 molar ratio for 1 hour at room temperature. To remove free dye and SFP, Mcm10-LD650 was diluted with 10% glycerol and 30 mM Tris-OAc pH 8.0 to reach a conductivity equal to 200 mM NaCl, loaded onto a column packed with 1 mL of sulphopropyl cation exchange resin (GE healthcare) and eluted with a 200-800 mM NaCl gradient in 10% glycerol, 30 mM Hepes pH 7.5, and 0.05% Tween 20. LD650-Mcm10 was eluted between 400 and 500 mM NaCl. The final labeling ratio was estimated to be ∼100% from the extinction coefficients of LD650 and Mcm10 (Figure S1B). Peak fractions were pooled, diluted to a conductivity equal to 250 mM NaCl with 10% glycerol, 30 mM Hepes pH 7.5, 4 mM DTT, and 40 μg/mL BSA, aliquoted, flash frozen, and stored at –80°C.

### Preparation of A488-RPA

*S. cerevisiae* RPA heterotrimer was purified as previously described (Henricksen et al., 1994). We used AlexaFluor488 (A488) NHS ester (Thermo Fisher) to nonspecifically label the primary amines of RPA. Preferential N-terminal labeling was achieved by labeling at low pH (7.0) for an NHS ester reaction (Selo et al., 1996). Labeling was performed with 50 mM Hepes pH 7.0, 150 mM NaCl, 1 mM DTT, and 0.25 mM EDTA. RPA was incubated with A488 dye at a 1:5 molar ratio for 1 hour at room temperature, and the reaction was quenched with 25 mM Tris-HCl pH 6.8 for 5 minutes. Excess dye was removed from RPA by buffer exchange with 20% glycerol, 30 mM Hepes pH 7.9, 150 mM NaCl, 1 mM DTT, and 0.25 mM EDTA. The final labeling ratio was estimated to be ∼90% from the extinction coefficients of A488 and RPA heterotrimer (Figure S1C). The final A488-RPA complex was aliquoted and stored at –80 °C.

### Preparation of Cy5-anti-Dig

Anti-digoxigenin antigen-binding fragments (Fab) (Roche) were resuspended in 0.1 M NaHCO_3_ pH 8.3 at 10 mg/mL. Cy5-NHS ester (GE Healthcare) was resuspended in DMSO. Dye was incubated with Fab at a 4:1 molar ratio for 1 hour at room temperature. The reaction was quenched with 25 mM Tris-HCl pH 6.8 for 5 minutes. Free dye was removed by a P-30 gel spin column (BioRad). The final labeling ratio was estimated to be ∼2.8:1. Cy5-anti-Dig was aliquoted, flash frozen, and stored at –80 °C.

### Preparation of biotinylated dsDNA

To create a terminally biotinylated double-stranded DNA template, the 12-base 5’ overhang on each end of genomic DNA from bacteriophage λ (48,502 bp; Roche) was filled in with a mixture of natural and biotinylated nucleotides by the exonuclease-deficient DNA polymerase I Klenow fragment (New England Biolabs). Reaction was conducted by incubating 10 nM λ-DNA, 33 μM each of dGTP/dATP/biotin-11-dUTP/biotin-14-dCTP (Thermo Fisher), and 5 U Klenow in 1× NEB2 buffer at 37 °C for 45 minutes, followed by heat inactivation for 20 minutes at 75 °C. DNA was then ethanol precipitated overnight at –20 °C in 2.5× volume cold ethanol and 300 mM sodium acetate pH 5.2. Precipitated DNA was recovered by centrifugation at 20,000× g for 15 minutes at 4 °C. After removing the supernatant, the pellet was air dried, resuspended in TE buffer (10 mM Tris HCl pH 8.0, 1 mM EDTA) and stored at 4 °C.

### Preparation of biotinylated ssDNA

Double-stranded λ DNA with one strand containing 4 biotins each at its 5’ and 3’ ends was obtained from LUMICKS (Amsterdam, Netherlands). A single dsDNA molecule was first captured between two beads, as assessed by a characteristic force-extension (*F*-*x*) curve (Smith et al., 1996). The non-biotinylated strand was then removed by overstretching the dsDNA at 0.1 μm/s under a low flow of 10 mM Tris HCl pH 8.0, based on a previously published protocol (Candelli et al., 2013). Establishment of a ssDNA tether was verified by its characteristic *F*-*x* curve (Figure S2) (Smith et al., 1996).

### Single-molecule data acquisition

Single-molecule experiments were performed at room temperature on a LUMICKS C-Trap instrument combining three-color confocal fluorescence microscopy with dual-trap optical tweezers (Hashemi Shabestari et al., 2017). A computer-controlled stage enabled rapid movement of the optical traps within a five-channel flow cell (Figure 1A). Laminar flow separated channels 1–3, which were used to form DNA tethers between 4.35-μm streptavidin-coated polystyrene beads (Spherotech) held in traps with a stiffness of 0.3 pN/nm. Under constant flow, a single bead was caught in each trap in channel 1. The traps were then quickly moved to channel 2 containing the biotinylated DNA of interest. By moving one trap against the direction of flow but toward the other trap, and vice versa, a DNA tether could be formed and detected via a change in the *F*-*x* curve. The traps were then moved to channel 3 containing only buffer, and the presence of a single DNA was verified by the *F*-*x* curve. Orthogonal channels 4 and 5 served as protein loading and/or experimental imaging chambers as described for each assay. Unless otherwise noted, flow was turned off during data acquisition. Force data was collected at 50 kHz. A488, Cy3 and LD650 fluorophores were excited by three laser lines at 488 nm, 532 nm and 638 nm, respectively. Kymographs were generated via a confocal line scan through the center of the two beads at 0.142 s/line (mode-switching data), 0.634 s/line (replication data), or 1.16 s/line (ssDNA data).

### Visualization of CMG on ssDNA

To investigate the ability of Cy3-CMG to load and translocate on ssDNA, optical traps tethering a single-stranded biotin-λ DNA (Figure S2) under a constant 5 pN tension were moved into channel 4 of the microfluidic flow cell (Figure 1A) containing 5 nM Cy3-CMG ± 10 nM Mcm10 in the following buffer: 25 mM Tris acetate pH 7.5, 5% glycerol, 40 μg/mL BSA, 3 mM DTT, 2 mM TCEP, 0.1 mM EDTA, 10 mM magnesium acetate, 50 mM potassium glutamate, and 1 mM ATP. In addition, the imaging buffer was supplemented with a triplet-state quenching cocktail (Dave et al., 2009) of 1 mM cyclooctatetraene (Sigma), 1 mM 4-nitrobenzyl alcohol (Sigma) and 1 mM Trolox (Sigma), as well as an oxygen scavenging system (Aitken et al., 2008) containing 10 nM protocatechuate-3,4-dioxygenase (Sigma) and 2.5 mM protocatechuic acid (Sigma). Following CMG loading, the tether was dragged to channel 5 that contained imaging buffer ± 1 mM ATP, supplemented with an ATP-regeneration system containing 60 μg/mL creatine phosphokinase (Sigma) and 20 mM phosphocreatine (Sigma).

### Detection of *in situ* DNA replication

To test whether Cy3-CMG can restart replication, optical traps tethering a double-stranded biotin-λ DNA under low tension were moved into channel 4 containing 30 nM Cy3-CMG, 60 nM Mcm10, and 1mM ATP. CMGM loading was promoted by stretching the tether to ∼65 pN (20 μm tether length, or ∼1.2 times its contour length). The CMGM-loaded tether was then dragged to channel 5 containing 5 mM ATP, and the additional *S. cerevisiae* proteins required for *in vitro* replication, purified as previously described (Georgescu et al., 2014; Lewis et al., 2017): 15 nM Pol α-primase, 15 nM Pol ε, 20 nM of Mrc1-Tof1-Csm3 heterotrimer, 5 nM RFC, and 25 nM PCNA. The imaging buffer described above was supplemented with 20 μM each of dATP/dCTP/dGTP/dTTP, and 100 μM each of rUTP/rCTP/rGTP. Upon lowering the force by decreasing the tether length to 15.1 μm (∼0.92 times its contour length) to promote strand reannealing and force collapse, directed motion of Cy3-CMG was observed (Figure 2D).

To confirm that this activity was due to replication, a similar experiment was performed in which optical traps tethering a double-stranded biotin-λ DNA under low tension were moved into channel 4 containing 30 nM Cy3-CMG, 60 nM Mcm10, 2 nM A488-RPA, 5 mM ATP, the additional *S. cerevisiae* proteins required for *in vitro* replication detailed above, as well as 20 μM each of dATP/dCTP/dGTP/dTTP, 100 μM each of rUTP/rCTP/rGTP, and 5 μM digoxigenin-11-dUTP (Roche). The tether was stretched to ∼65 pN in order to create single-stranded regions and promote replisome assembly at the fork, determined by visualization of Cy3-CMG and A488-RPA. Following replisome loading, the tether was held at a constant low force for 5–10 minutes (Figure S6A). Replication was assessed by detection of Dig-dUTP incorporation by dragging the tether to channel 5 containing imaging buffer supplemented with 50 nM Cy5-anti-Dig Fab (Figure S6B). Bulk analysis of replication in the presence of Dig-dUTP revealed severe inhibition of replisome speed (Figure S6C), accounting for the lack of CMGM movement in the single-molecule Dig-dUTP replication experiments.

### Observation of CMG mode-switching

Optical traps tethering a double-stranded biotin-λ DNA under a constant 10 pN tension were moved into channel 4 containing 30 nM Cy3-CMG ± 60 nM Mcm10 in imaging buffer supplemented with 5 mM ATP. The tether was stretched to ∼65 pN in order to create ssDNA regions and promote CMG loading at the fork. Following the visualization of CMG loading, the tension was lowered to a constant force of 10 pN. Repeated cycles of this sequential force protocol were conducted to increase throughput. This assay was also performed as two-color (2 nM A488-RPA + 30 nM Cy3-CMG; or 30 nM Cy3-CMG + 60 nM LD650-Mcm10) experiments.

### Bulk helicase unwinding assay

Circular M13 bacteriophage ssDNA (6.4 kbp; New England Biolabs) was annealed to a 5’- ^32^P labeled oligo containing a 5’-dT_60_ flap and a 35-nt sequence at its 3’ end that is complementary to M13 (Figure S5). 0.5 nM of this substrate was incubated with 20 nM CMG in the presence or absence of 40 nM Mcm10 for 2 minutes. Reactions were initiated with 1 mM ATP, followed by an addition of 20 μM unlabeled flap oligo 2 minutes after initiation to prevent unwound radiolabeled oligo from reannealing to the M13 substrate. Aliquots were stopped with a final concentration of 1% SDS and 20 mM EDTA and flash frozen in liquid nitrogen. All reactions took place at 30 °C and in 5% glycerol, 40 mg/mL BSA, 5 mM TCEP, 10 mM magnesium sulfate, 25 mM KCl, 25 mM Tris acetate pH 7.5, and 0.1 mM EDTA. Flap displacement was resolved by 10% PAGE, exposed to a phosphorimager screen, and imaged with Typhoon 9500 (GE healthcare).

### Bulk replication assay

Unprimed, nucleotide-biased 3.2-kb forked DNA substrates that lack dC on the leading strand and dG on the lagging strand were used such that incorporation of ^32^P-α-dCTP reported on leading-strand replication only (Schauer et al., 2017). DNA substrate (1.5 nM) was incubated with 40 nM CMG and 60 nM Mcm10 in the presence or absence of 5 μM Dig-dUTP for 10 minutes at room temperature. An enzyme mix containing 15 nM Pol α, 15 nM Pol ε, 30 nM MTC, 5 nM RFC, 25 nM PCNA, 20 μM dATP, and 20 μM dCTP was added for 2 minutes at room temperature. The reaction was initialized with 5 mM ATP, 20 μM dGTP, 20 μM dTTP, and 125 nCi/μL ^32^P-α-dCTP, and aliquots were stopped with a final concentration of 1% SDS and 20 mM EDTA. Reactions were run on 1.2% (wt/vol) alkaline agarose for 17 hours at 35 V, backed with DE81 paper, and dried by compression. The gel was exposed to a phosphorimager screen and imaged with Typhoon 9500 (GE Healthcare). Reaction volumes were 20 μL and took place at room temperature and in 5% glycerol, 80 μg/mL BSA, 5 mM TCEP, 10 mM magnesium acetate, 50 mM potassium glutamate, 25 mM Tris acetate pH 7.5, and 0.1 mM EDTA.

### Data analysis

Single-molecule force and fluorescence data were analyzed using custom software provided by LUMICKS. The loading efficiency of CMG on ssDNA was determined per tether by dividing the number of CMG complexes stably bound to ssDNA by the incubation time (∼5 minutes in the absence of Mcm10 and ∼30 seconds in its presence) in the CMG channel (channel 4) and the CMG concentration (5 nM). Only stably bound CMG molecules were considered, defined as those that survived dragging from channel 4 to channel 5. 62 out of 92 bound CMGs (67%) displayed unidirectional translocation and did so at an approximately constant speed (Figure 1B), enabling rate estimation by dividing the distance traveled by the duration of each trajectory. Mean square displacement (MSD) analysis of CMG diffusion was performed as follows. Segments of kymographs containing diffusive CMGs were cropped in Fiji (Schindelin et al., 2012). The KymographClear Fiji plugin and the standalone KymographDirect software (Mangeol et al., 2016) were used to extract position versus time information from each diffusing CMG trajectory. MSD calculations were performed in MATLAB using the @msdanalyzer class (https://github.com/tinevez/msdanalyzer) (Tarantino et al., 2014) with custom modifications. 35 out of 48 analyzed trajectories (73%) displayed a linear MSD profile for ∆*t* from 0 to 25% of the length of the trajectory [R^2^ = 0.96 ± 0.05 (mean ± SD) for linear regression]. A diffusion coefficient (*D*) was obtained for each CMG trajectory that fits this criterion (Figure S8). Unless otherwise noted, errors reported in this study represent the approximated standard error of the mean determined from 100,000 bootstrapped samples for each dataset. *P*-values reported were determined from two-tailed two-sample *t*-tests.

## Supplementary Figures

**Figure S1.**
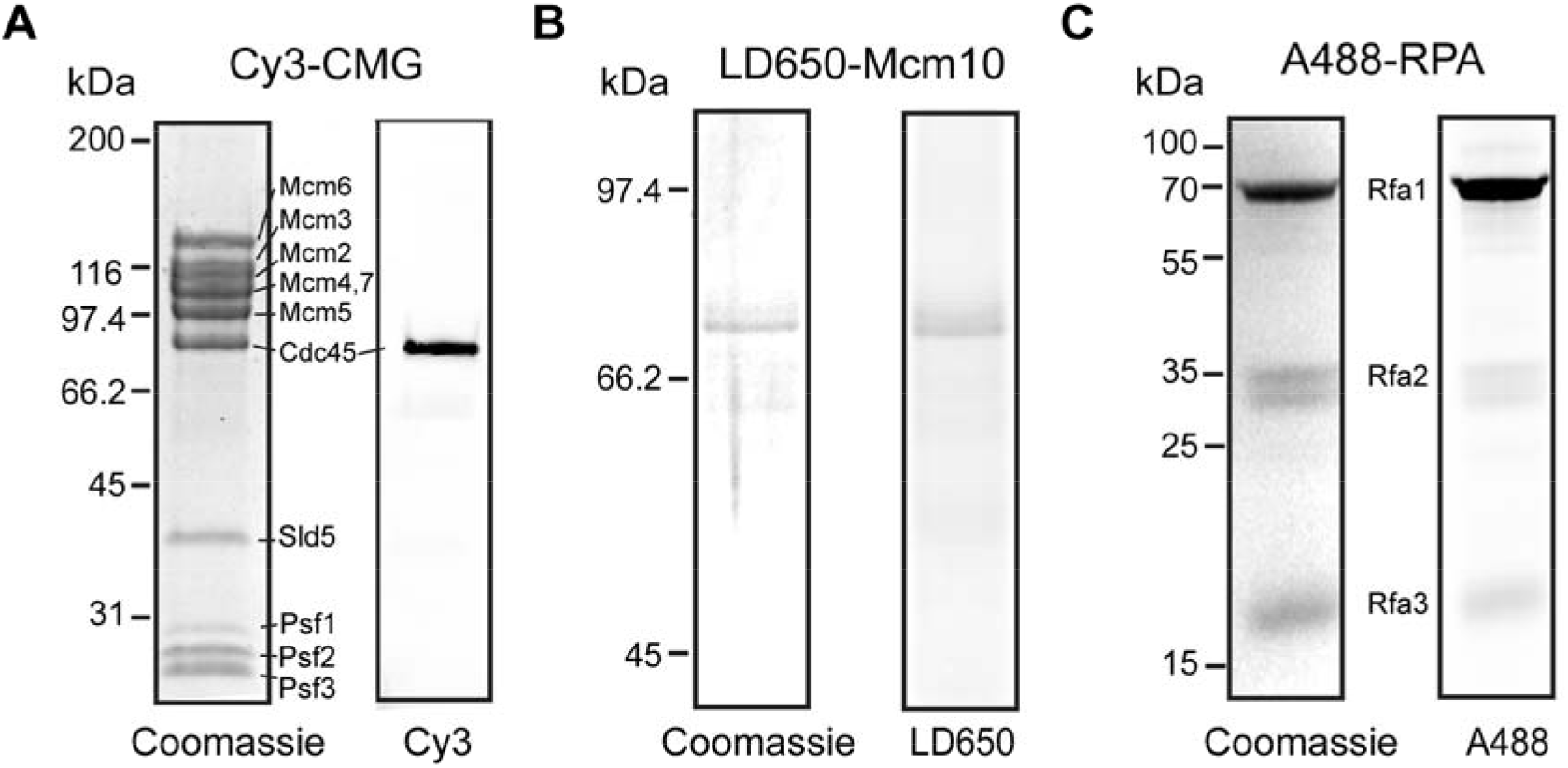
Protein purification and fluorescent labeling. The purity of Cy3-CMG (**A**), LD650-Mcm10 (**B**), and A488-RPA (**C**) was analyzed by SDS-PAGE via Coomassie Blue staining (Left) and fluorescence scan (Right). See Materials and Methods for detailed purification and labeling procedures.

**Figure S2.**
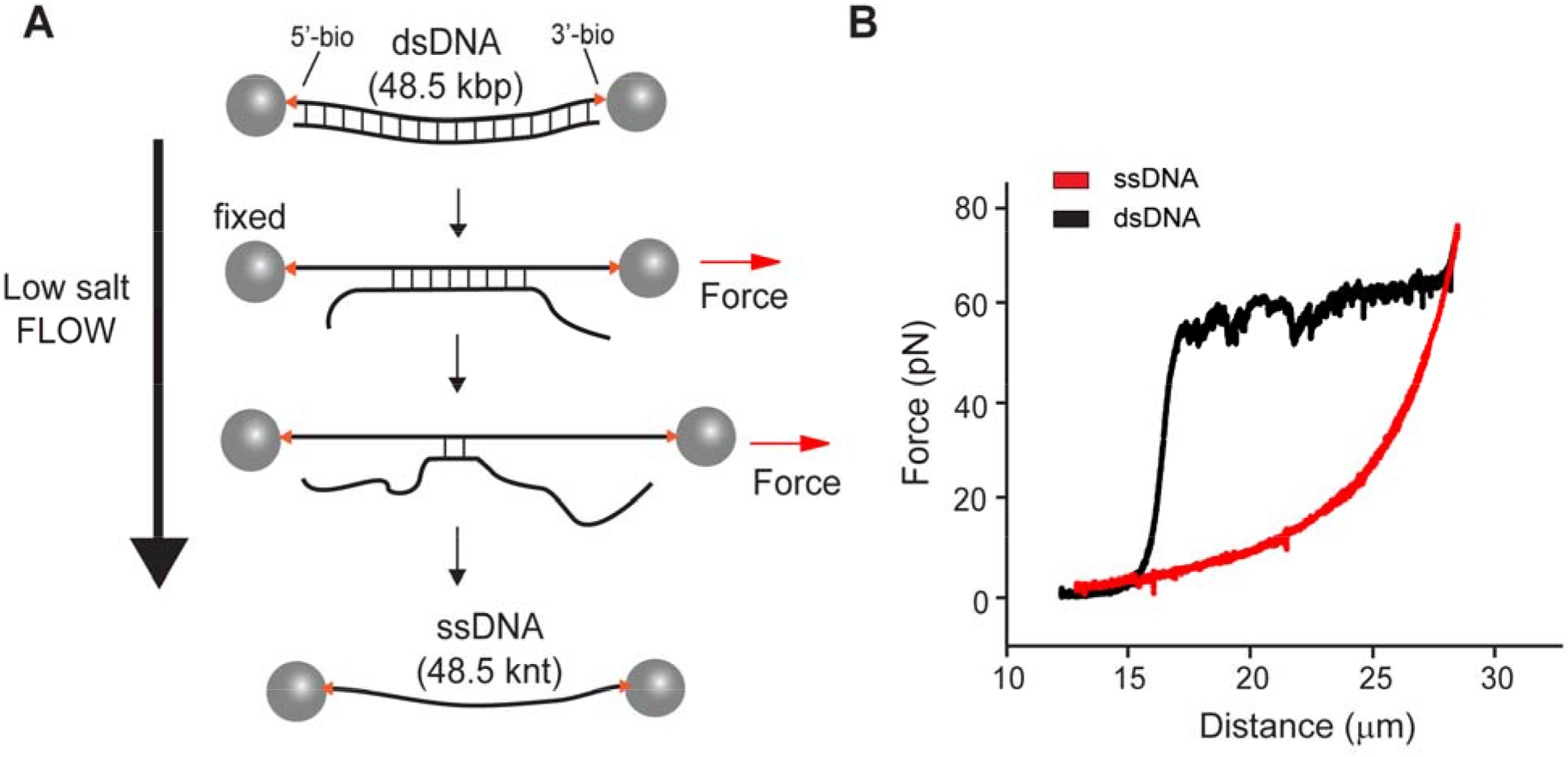
Generation of single-stranded DNA tethers. (**A**) Schematic illustrating *in situ* formation of a ssDNA. Two optically trapped streptavidin-coated beads were used to capture a single dsDNA biotinylated on both ends of one strand. The non-biotinylated strand was then removed by force-induced melting of the duplex, aided by low flow in a buffer lacking salt. (**B**) Force-extension curve indicating the transition from dsDNA (black) to ssDNA (red).

**Figure S3.**
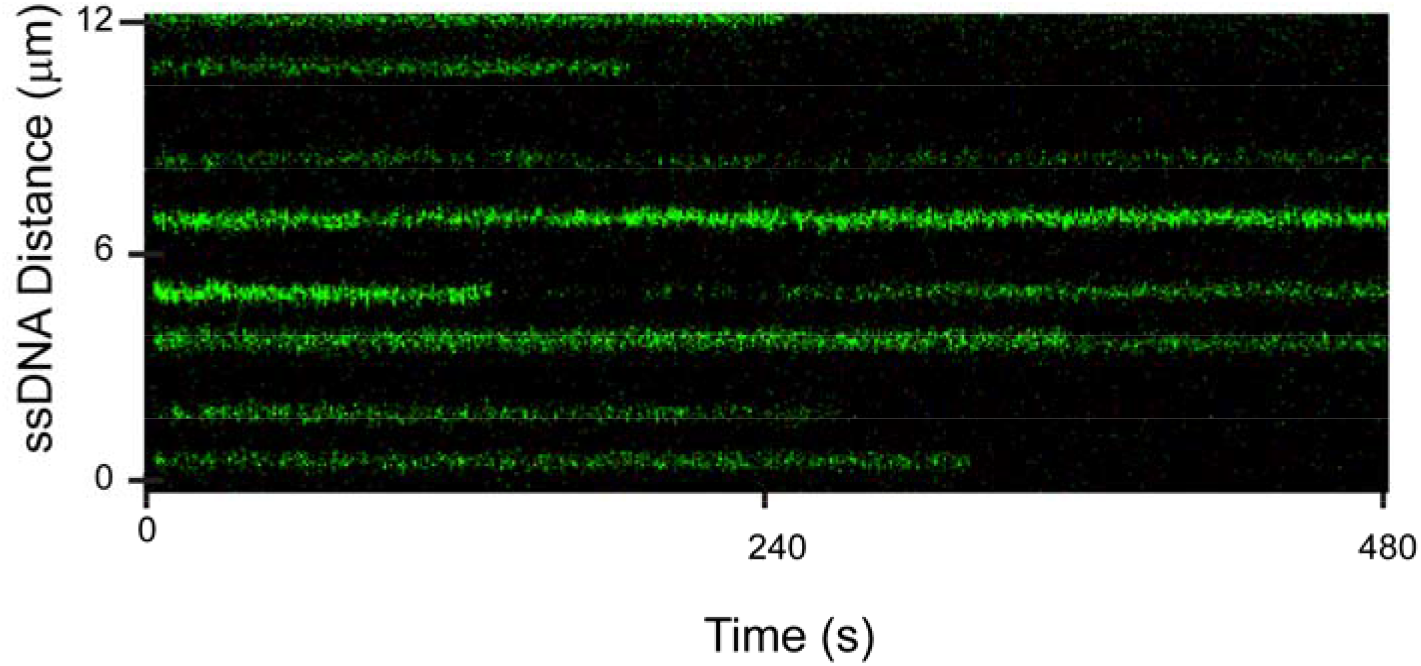
Directed motion of CMG on ssDNA does not occur without ATP. Representative kymograph of Cy3-CMG (green) and Mcm10 loaded onto ssDNA under 5 pN tension and imaged in the absence of ATP. The directed motion that normally occurs in the presence of ATP (Figure 1B) was not observed (0 out of 43 CMGs).

**Figure S4.**
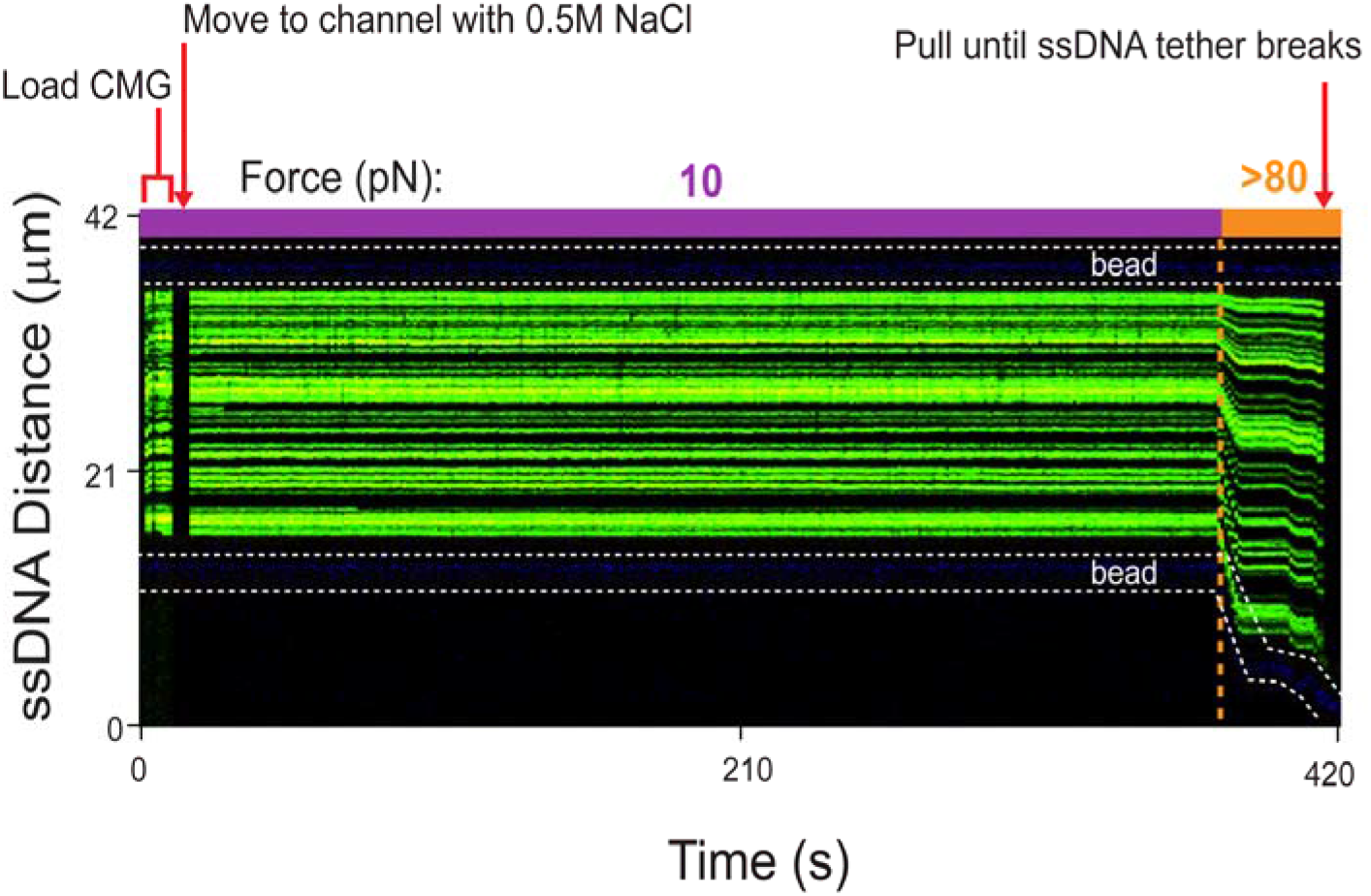
CMGM is stable on ssDNA at high salt and under high tether tension. Shown is a kymograph in which Cy3-CMG (green) and Mcm10 was first loaded onto ssDNA at low salt (50 mM potassium glutamate) and low force (10 pN) in the absence of ATP. The tether was then dragged to a new channel containing high salt (0.5 M NaCl + 50 mM potassium glutamate) followed by an increase of force applied to the DNA to >80 pN. In this example, the ssDNA tether was incubated with an elevated CMGM concentration compared to the experiments described in the main text (e.g., in Figure 1B) in order to overload the tether with CMGM. The ability of CMG to withstand hydrodynamic dragging, high salt and high tension in the tether collectively suggests that it encircles ssDNA.

**Figure S5.**
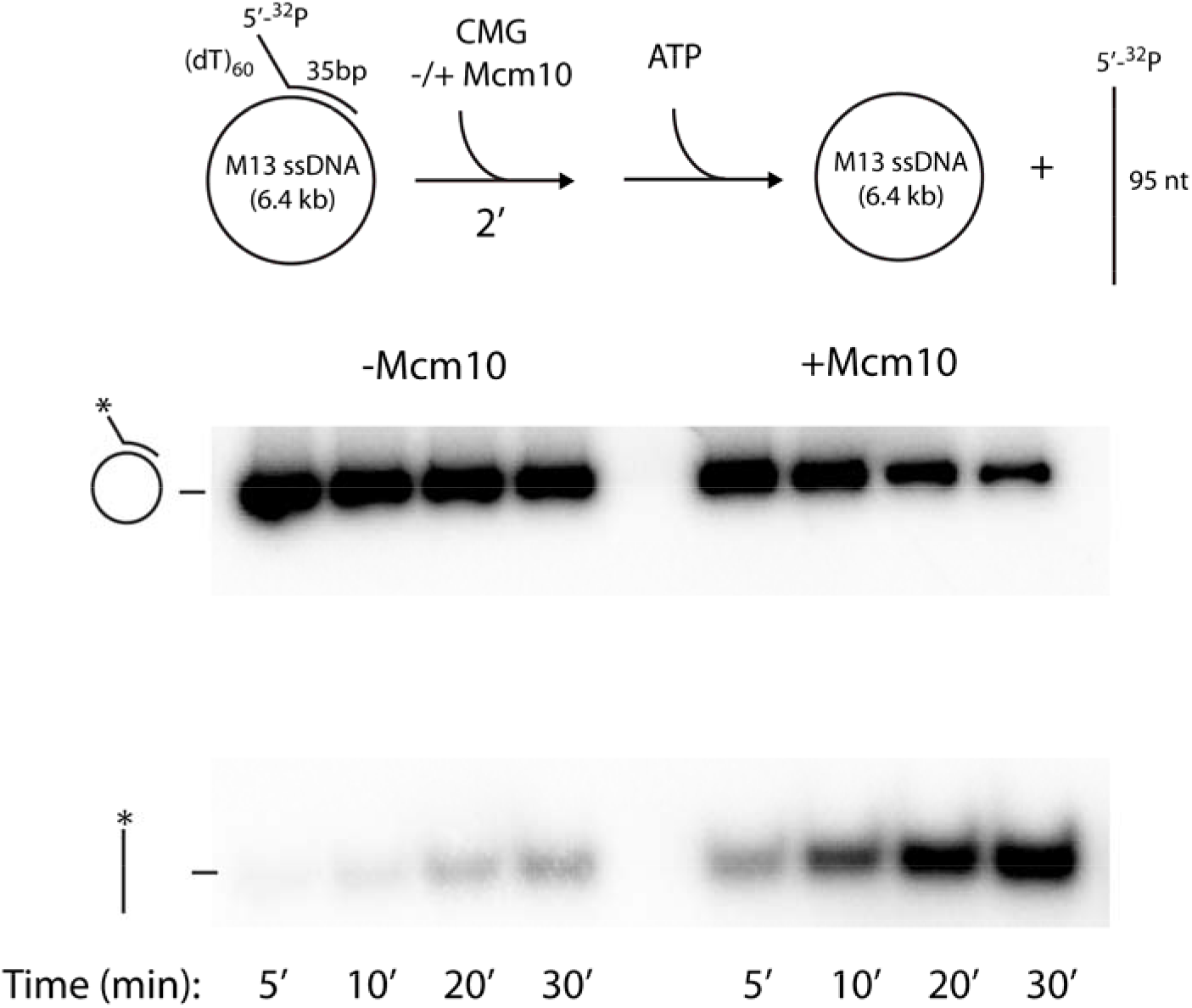
Displacement of a 5’ -^32^P oligonucleotide from circular M13 ssDNA. Reactions were performed in the absence or presence of Mcm10. CMG and CMGM must encircle the circular strand in order to unwind the 5’- ^32^P oligonucleotide. Reactions were stopped at the indicated timepoints.

**Figure S6.**
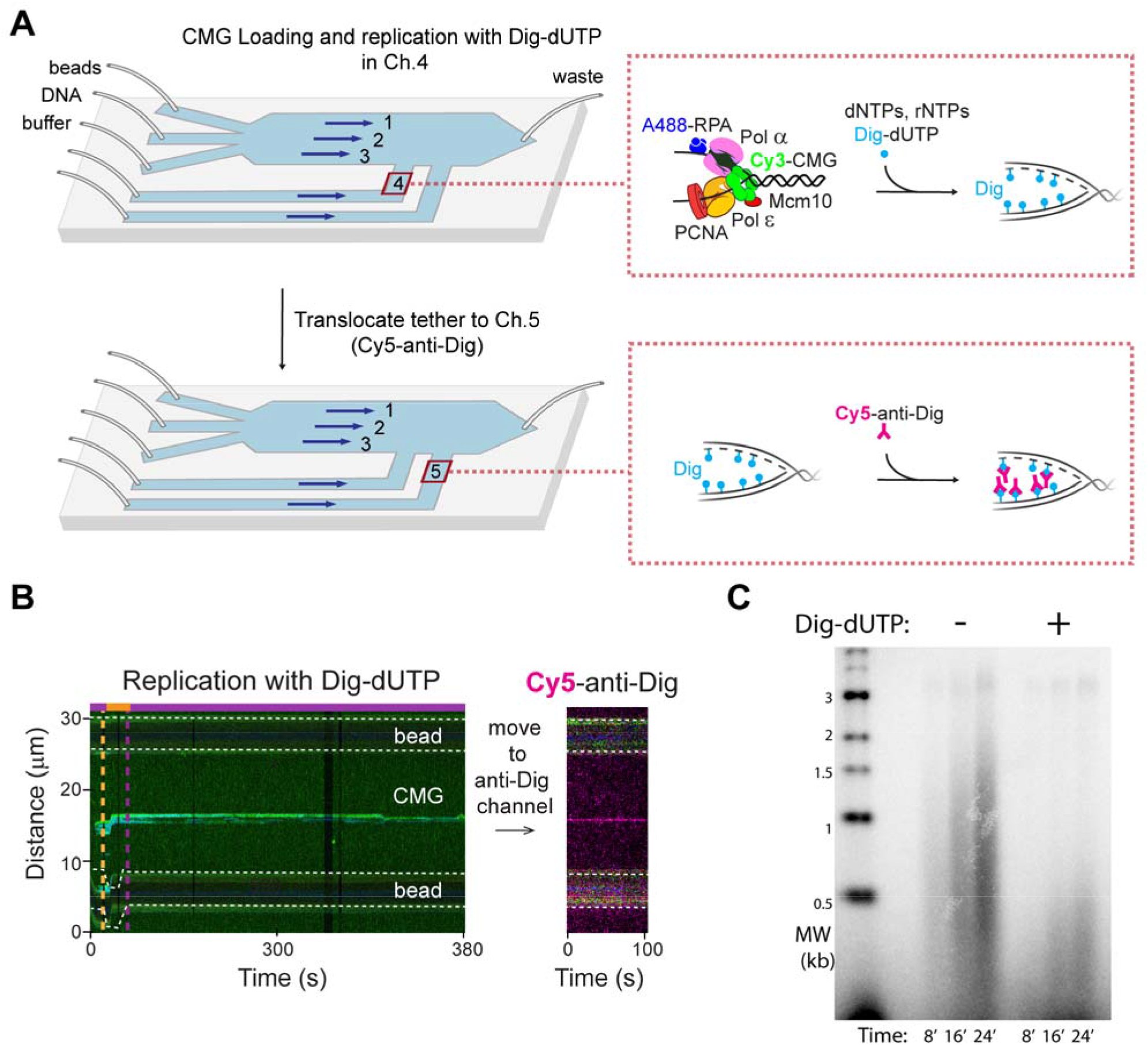
CMGM loading at the fork leads to active DNA replication. (**A**) Schematic of the experimental setup for detection of Dig-dUTP incorporation. (Top) Cy3-CMGM and other unlabeled replisome components were loaded at a DNA fork initially marked by A488-RPA and incubated with a full set of nucleotides and Dig-dUTP. (Bottom) The post-replication assembly was moved to a separate channel containing Cy5-anti-Dig for staining nascent DNA. (**B**) A representative kymograph of *in situ* DNA replication using Dig-dUTP. The nascent DNA tract (magenta) was observed at the same location as CMGM. (**C**) Dig-dUTP inhibits replication. Replication of 3.2-kb forked DNA substrate was stopped at the indicated timepoints (see Materials and Methods).

**Figure S7.**
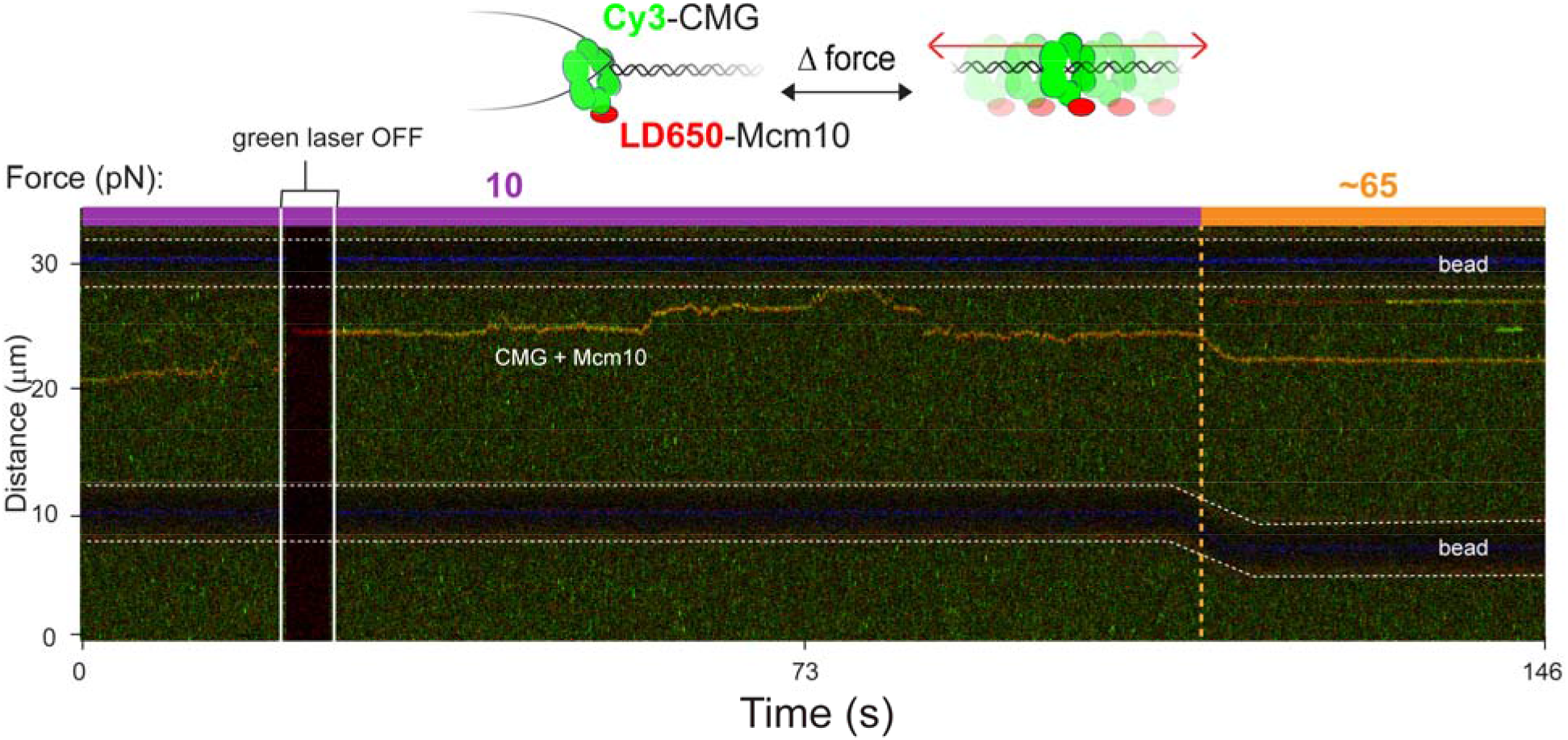
Diffusion of the CMGM complex on dsDNA. Experiments performed with Cy3-CMG (green) and LD650-Mcm10 (red) demonstrate that Mcm10 travels with CMG in the diffusive mode. The green laser was occasionally turned off to confirm the red fluorescence signal from Mcm10.

**Figure S8.**
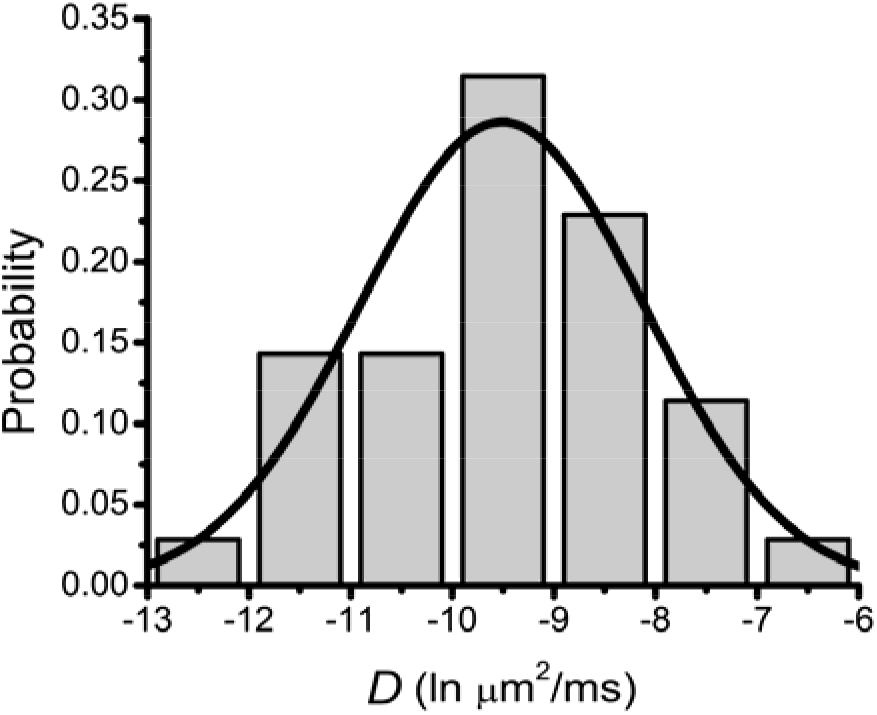
Mean square displacement (MSD) analysis for CMGM diffusion. Diffusion coefficients (*D*) were estimated by linear regression of the MSD plots from diffusive CMGM trajectories (Figures 3E and 3F; *N* = 35). The histogram was well fit to a lognormal distribution with *D* = 0.20 ± 0.052 µm^2^/s or 1.81 ± 0.48 kbp^2^/s (mean ± SEM).

**Figure S9.**
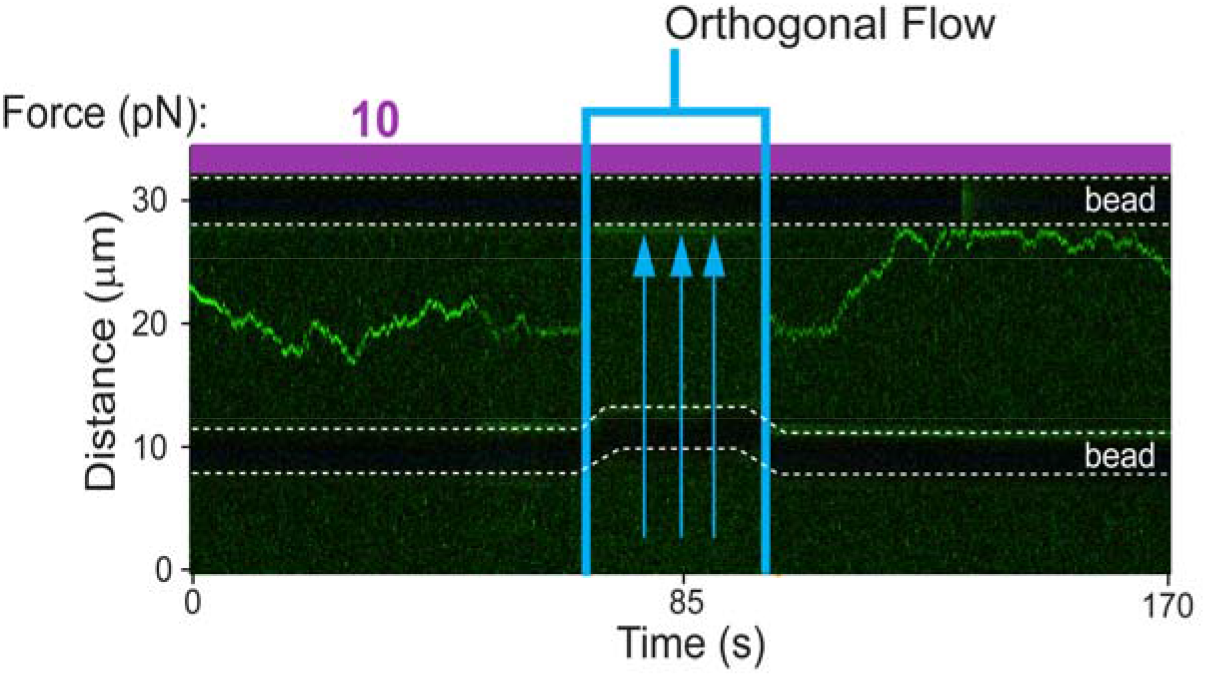
Robust diffusive behavior of CMGM on dsDNA. CMGM, while in the dsDNA diffusive mode, persisted on dsDNA under hydrodynamic force generated by an orthogonal flow, suggesting that CMGM is topologically linked to dsDNA during rapid diffusion. The tether temporarily went out of the imaging plane when the flow was applied.

**Figure S10.**
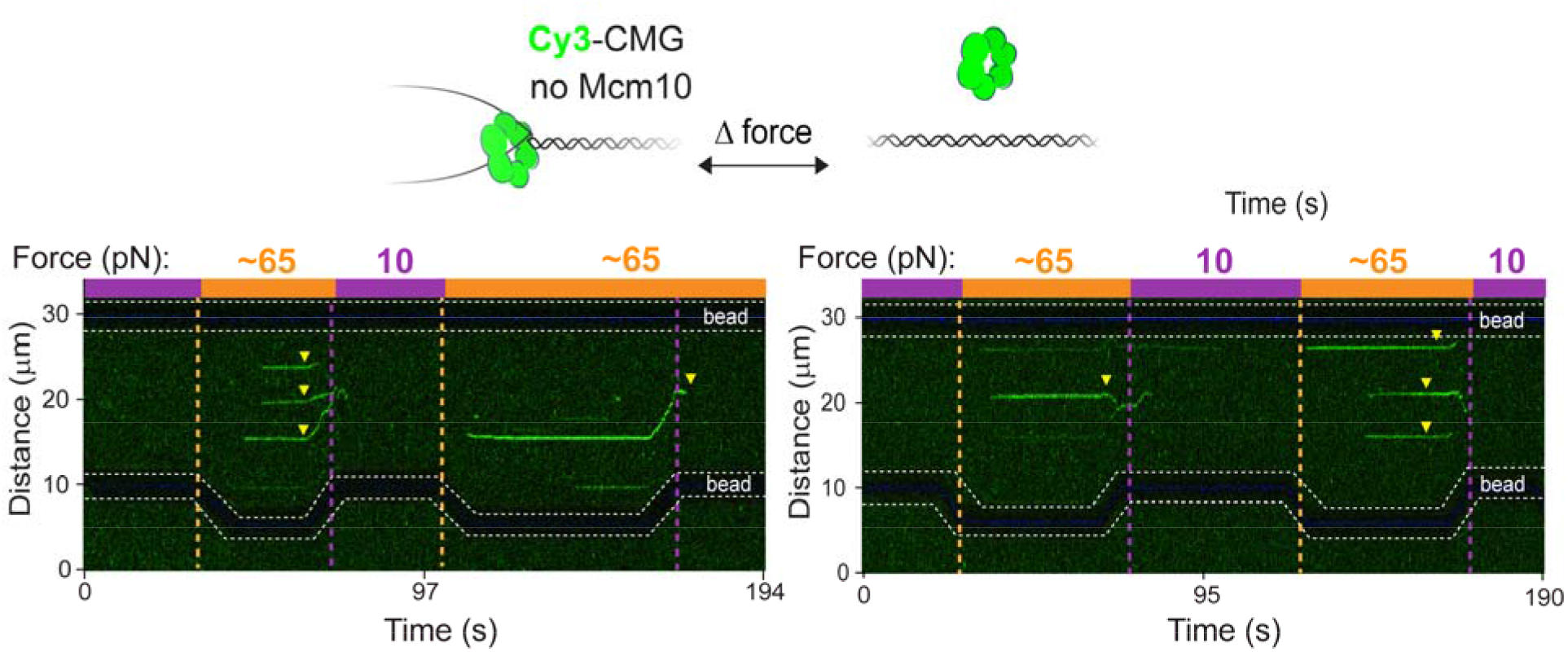
Mcm10 stabilizes CMG in the diffusive mode. Representative kymographs show that, in the absence of Mcm10, CMG (green) predominantly dissociates (triangle) from DNA following fork collapse.

